# Ultrasound-led stratification of carpal tunnel syndrome reveals structure–function mismatch

**DOI:** 10.64898/2026.05.09.723950

**Authors:** Jishizhan Chen, Daqian Shi, Jiaqi Su, Xiaohai Huang, Yun Qian

## Abstract

The severity stratification of carpal tunnel syndrome (CTS) relies on ultrasound morphological markers and electromyography. However, it remains unclear how structural imaging can reliably infer functional impairment. Clarifying the structure–function relationship is critical for efficient diagnostic pathways. A retrospective cohort of 55 patients with symptoms related to CTS was analyzed at the Shanghai Sixth People’s Hospital. All patients were subjected to ultrasound and EMG. 72.7% cases were diagnosed with CTS with a female predominance and equal left–right involvement. Random-forest classifiers were trained using surrogate splits, and performance was evaluated using predictions outside the bag. A full-feature model (34 candidate variables) was compared against a simplified model (8 core variables) capturing the core morphological and electrophysiological features. A residual-based framework was then used to characterize the structure–function mismatch within severity grades (1a–3c). The simplified model improved discriminative performance compared to the full-feature model (AUC 0.789 to 0.824). The simplified model achieved an overall accuracy of 77.3%. Analysis of predicted probability distributions and 10-bin calibration curves indicated stable and clinically interpretable risk estimation in most probability ranges. Permutation-based importance analysis confirmed that both ultrasound and electrophysiological features contributed substantively to prediction. Residual-based grading further revealed structure– function heterogeneity within each main severity grade. CTS severity can be stratified using a limited set of complementary morphological and electrophysiological features. Structure–function mismatch supports an imaging-led initial screening, with electrophysiology reserved for selected patients.

## 1. Introduction

Carpal tunnel syndrome (CTS) is the most common entrapment neuropathy of the upper limb and a major cause of pain and work disability[4]. Clinical management of CTS depends not only on establishing the diagnosis, but also on accurately assessing disease severity to guide treatment decisions. CTS diagnosis in the past relied mainly on characteristic nocturnal paresthesia and physical tests but showed limited specificity and reproducibility[5]. The subsequent introduction of ultrasound and electromyography enabled quantitative assessment of median nerve morphology and dysfunction, becoming a cornerstone of CTS diagnosis. According to the 2025 evidence-based clinical practice guideline from the American Academy of Orthopaedic Surgeons[10], the diagnosis of CTS remains primarily clinical, based on characteristic symptoms in the median nerve distribution combined with physical examinations. Ultrasound and electromyography are recommended as objective approaches to assess disease severity but are not recommended as standalone diagnostic criteria or gold standards.

Reliable severity assessment remains a real-world challenge due to two main factors: poor data quality (incompleteness, redundancy, and measurement variability) and the complex relationship between structural compression and functional nerve impairment[7]. Ultrasound provides a non-invasive and accessible assessment of median nerve morphology[7]. Morphological markers such as nerve swelling, flattening, and changes in surrounding structures have demonstrated diagnostic value. However, it does not directly reflect the nerve function. Electromyography assesses functional impairment of the median nerve and are often regarded as a supplementary approach for CTS diagnosis[11]. However, it is invasive and uncomfortable, which may reduce patient compliance and result in incomplete testing[6]. Importantly, the relationship between morphological abnormalities and electrophysiological impairment is not straightforward. While severe CTS often presents with consistent morphological and electrophysiological changes, patients with mild or moderate CTS exhibit ambiguous findings[12]. Electrophysiological impairment may be present in the absence of significant morphological abnormalities, vice versa. Despite expert consensus that combining ultrasonography and electrophysiology provides more information[8], the effectiveness of these two techniques in CTS and the importance of various assessment features remain unclear[9]. Therefore, we hypothesize that excessive features in both ultrasound and electromyography introduce unnecessary noise, screening core features can improve the interpretability without additional examinations.

Clarifying the structure–function relationship in CTS is critical for evidence-based severity stratification. In this study, we use both real-world ultrasound and electrophysiological data with heterogeneity and incompleteness to construct both full-feature (34 variables) and simplified (eight core variables) models to predict CTS severity. We proved that CTS severity can be stratified using a limited set of complementary features, and structure–function mismatch reveals clinically meaningful subgroups.

## 2. Experiments

### 2.1. Study cohort and data acquisition

This retrospective study included 55 patients presenting with symptoms of CTS at Shanghai Sixth People’s Hospital (processed dataset available via Zenodo at https://doi.org/10.5281/zenodo.18349319). All subjects un-derwent both wrist ultrasound examination and electrophys-iological tests using routine clinical protocols. Each wrist was treated as an independent sample, resulting in bilateral data where available. All ultrasound images were stored in DICOM format, and text reports were in JPEG format. All electrophysiological text reports were stored in JPEG format. Clinical diagnosis of CTS was established by experienced clinicians based on ultrasound and electrophysiological reports and standard clinical criteria.

### 2.2. Feature definition, selection, and preprocessing

A total of 34 candidate variables were selected from both ultrasound-derived morphological features and electrophysiological measurements. The ultrasound examinations assessed median nerve morphology at anatomical levels of mid forearm, pisiform, and hamate. Reported morphological features included transverse and anteroposterior diameters, cross-sectional area, and carpal tunnel narrowest point. The electrophysiological tests comprised median nerve conduction studies, including conduction velocity, distal latency, and nerve action potential amplitudes (Table 1). Quantitative variables from all reports in JPEG format were extracted using RZCR[13] and CharFormer[3], followed by structured formatting into tabular form. Extracted values were manually verified to correct recognition errors. Given the inherent heterogeneity and incompleteness of clinical data, samples were excluded if less than 30% of candidate variables were available. No imputation was performed prior to model training. All 34 candidate variables were used to train the full-feature model.

**Table 1.**
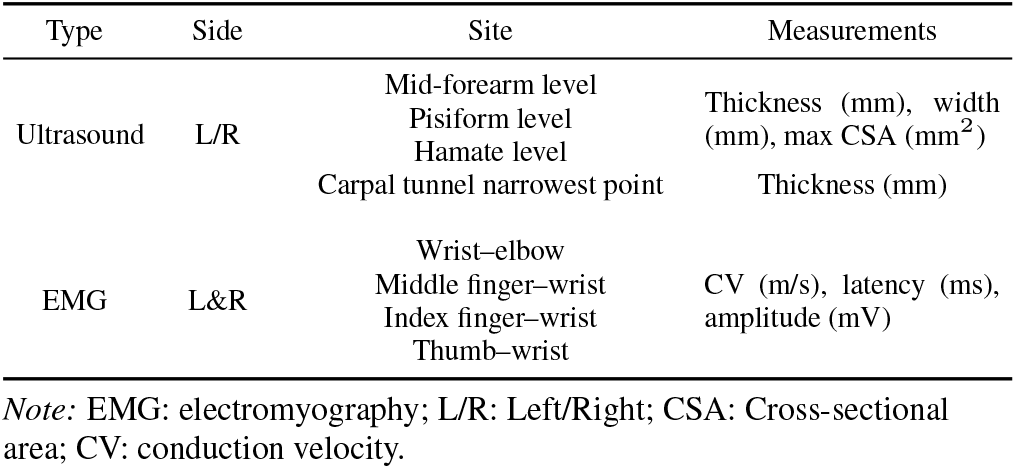
Summary of full candidate variables from ultrasound and electromyography.

### 2.3. Feature importance analysis

To assess whether CTS identification and severity stratification could be achieved with a simplified model, feature importance analysis was performed. Cross-validated permutation importance (area under the curve (AUC), average precision, F1-score, accuracy, and Brier Score) with model-based measures (mean decrease in impurity and split counts) were utilized to capture complementary morphological and electrophysiological features. As a result, a subset of eight core variables was determined to train the simplified model (Table 2).

**Table 2.** Summary of eight core variables determined by importance analysis.

| Modality | Side | Variable |
| --- | --- | --- |
| Ultrasound | L/R | CSA at the pisiform level (mm<sup>2</sup>) |
|  |  | CSA at the mid-forearm level (mm<sup>2</sup>) |
| EMG | L&R | Wrist–elbow amplitude (mV) |
|  |  | Middle finger CV (m/s) |
|  |  | Middle finger latency (ms) |

Note: EMG: electromyography; L/R: Left/Right; CSA: Cross-sectional area; CV: conduction velocity.

### 2.4. Development and evaluation of full-feature and simplified models

Supervised classification full-feature (34 candidate variables) and simplified (8 core variables) models were implemented using random forests (MATLAB TreeBagger, 300 decision trees). Random forests were selected due to their robustness to small sample sizes, multicollinearity, and partial missingness. Surrogate splits were enabled to allow tree-based decisions when primary splitting variables were unavailable.

Model performance was evaluated using out-of-bag (OOB) predictions, providing an internal and approximately unbiased estimate of generalisation performance without requiring an explicit hold-out test set. Discriminative performance was quantified using the AUC, sensitivity, specificity, overall accuracy, and the Kolmogorov–Smirnov statistic. Operating points were selected based on maximising the Youden index unless otherwise stated. To assess probabilistic reliability, predicted probability distributions were examined and calibration curves were constructed using decile (10-bin) grouping of predicted risk. Observed and predicted event rates across probability ranges were compared.

### 2.5. Residual-based grading and structure–function mismatch

To characterise structure–function mismatch within CTS severity grades, a residual-based grading framework was applied using the simplified feature set. The model was constructed by regressing nerve conduction velocity on median nerve cross-sectional area at the pisiform level and the wrist-to-forearm ratio. Left and right wrists were combined to increase sample size. For each wrist, a residual was calculated as the difference between observed and model-predicted conduction velocity. Positive residuals indicate relatively preserved nerve function given the degree of morphological abnormality, whereas negative residuals indicate disproportionately impaired function relative to morphology.

Main CTS severity grades (0–3) were first assigned using predefined ultrasound and electrophysiological parameters. Residual-based subgrading (a–c) was then performed within each main grade by ranking residuals and partitioning them into tertiles. These subgrades represent better than expected, approximately consistent, and worse than expected functional status within the same main grade.

### 2.6. Statistical analysis

All analyses were conducted using MATLAB (MathWorks, Natick, MA, US) and Microsoft Excel (Microsoft Corporation, Redmond, WA, USA). Continuous variables are reported as mean ± standard deviation unless otherwise stated.

## 3. Experiment Results

### 3.1. Cohort characteristics and data completeness

A total of 55 patients presenting with CTS symptoms were included. Overall, 72.7% of wrists were clinically diagnosed as CTS-positive, with a female predominance (15% male, 85% female) and equal left–right involvement (Figure 1a). Age distributions overlapped between male and female CTS patients, with peak at 52.0 years old and 53.2 years old, respectively (Figure 1b).

**Figure 1.**
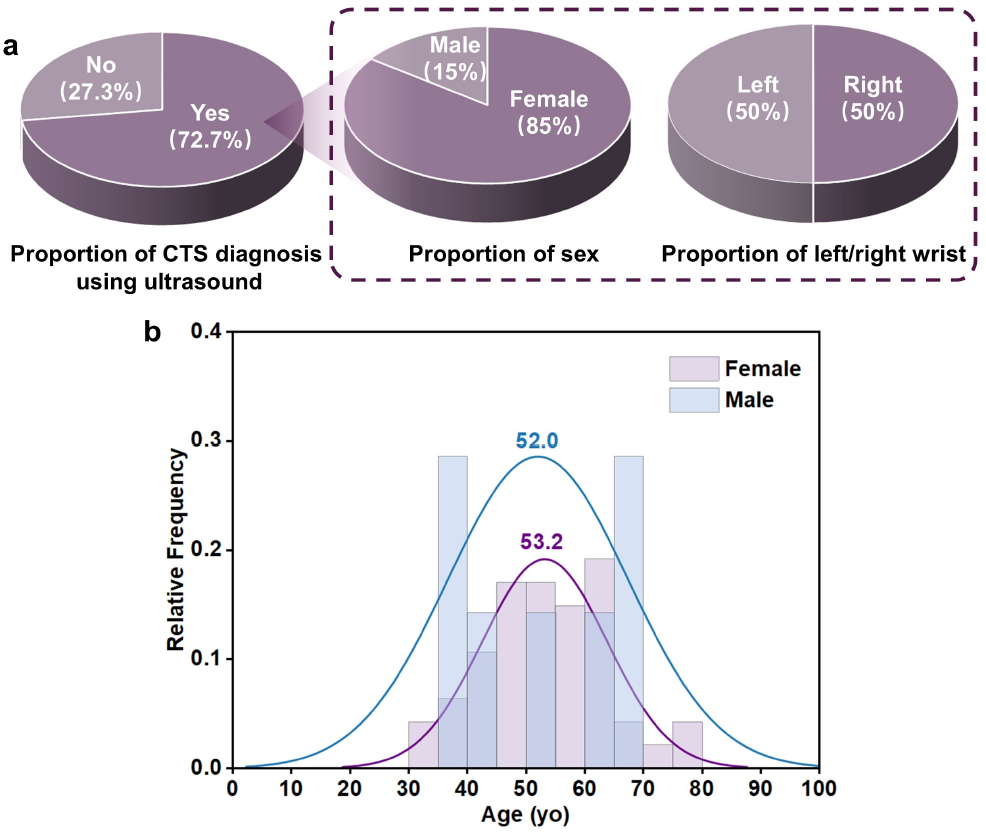
Cohort characteristics. (a) Proportion of CTS diagnosis, sex, and wrist side; (b) Age distribution of male and female CTS patients.

Across the dataset, 34 candidate variables were selected from ultrasound and electrophysiological reports. Feature completeness varied substantially across variables and samples. Applying a predefined inclusion criterion of ≥30% feature availability, 80% of wrist-level samples (84/105) were retained for modelling. Incompleteness affected both ultrasound and electrophysiological variables, reflecting the significant heterogeneity of clinical data.

### 3.2. Performance of the full-feature model

Using all 34 candidate variables, the random forest classifier achieved discriminative performance based on OOB predictions (Figure 2). Receiver operating characteristic (ROC) analysis yielded an AUC of 0.789. The optimal operating point at a probability threshold of 0.806 resulted in a sensitivity of 67.2% and a specificity of 88.2%, corresponding to an overall accuracy of 71.4%. The Kolmogorov–Smirnov statistic reached 0.554, indicating strong separation between CTS-positive and CTS-negative probability distributions.

**Figure 2.**
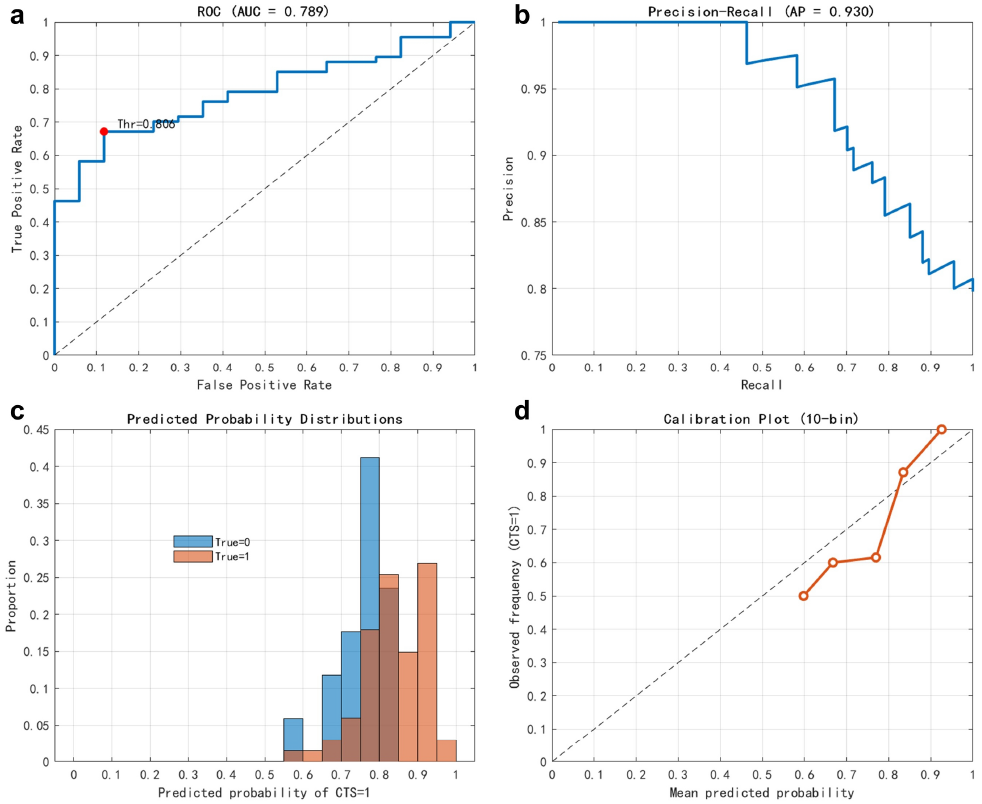
Performance of the full-feature model. (a) ROC curve with AUC = 0.789; (b) Precision–recall curve with AP = 0.930; (c) Distribution of predicted probabilities stratified by true class (CTS= 1, no CTS = 0); (d) Calibration plot (10-bin) comparing mean predicted probabilities with observed CTS frequencies.

The corresponding confusion matrix comprised 45 true positives, 15 true negatives, 22 false negatives, and 2 false positives, reflecting a conservative operating profile prioritising specificity. Precision–recall analysis showed robust performance under class imbalance, with an average precision (AP) of 0.930. Precision remained high across low-to-moderate recall levels and declined at higher recall values, consistent with increasing overlap among intermediate-risk cases.

Predicted probability distributions demonstrated enrichment of CTS-positive samples at higher probability ranges, while CTS-negative samples clustered at lower probabilities. Calibration analysis using 10-bin (decile) grouping showed good agreement between predicted and observed CTS prevalence across most probability ranges, with minor deviations in intermediate bins.

Permutation-based importance analysis identified multiple influential variables crossing both modalities. Ultrasound features and electrophysiological measurements each contributed substantively to prediction. This further proved the complementary nature of structural and functional assessment in CTS. No single feature or modality dominated model performance, indicating that CTS severity cannot be reliably inferred from morphology or electrophysiology alone. This finding aligns with clinical experience that structural compression and functional impairment in CTS do not progress in a synchronous manner[1].

### 3.3. Performance of the simplified model

Restricting the model to eight pre-specified core variables resulted in improved discriminative performance compared with the full-feature model (Figure 3). The simplified model yielded an AUC of 0.824, increased by 0.035, compared to the full-feature model. This indicated enhanced class separation despite feature reduction.

**Figure 3.**
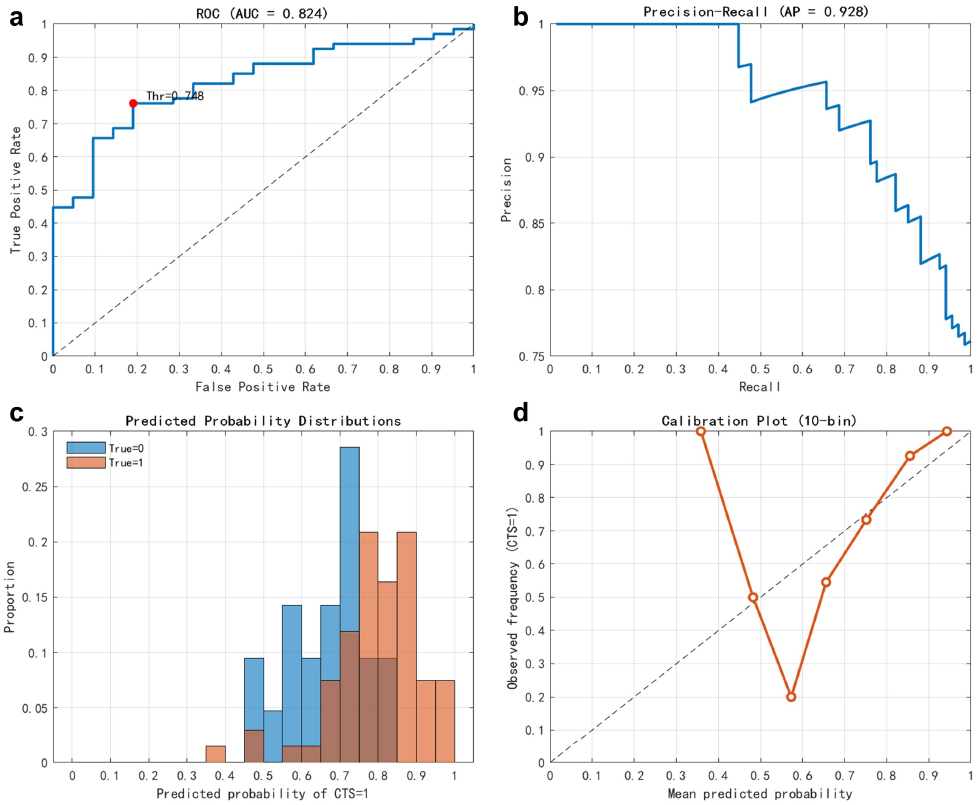
Performance of the simplified model. (a) ROC curve with AUC = 0.824; (b) Precision–recall curve with AP = 0.928; (c) Distribution of predicted probabilities stratified by true class (CTS = 1, no CTS = 0); (d) Calibration plot (10-bin) comparing mean predicted probabilities with observed CTS frequencies.

At the optimal operating point at a probability threshold of 0.748, the simplified model achieved a sensitivity of 76.1% and a specificity of 81.0%, corresponding to an overall accuracy of 77.3%. The Kolmogorov–Smirnov statistic increased to 0.571, reflecting improved separation between cumulative probability distributions relative to the full model. The corresponding confusion matrix comprised 51 true positives, 17 true negatives, 16 false negatives, and 4 false positives, indicating improved recall of CTS-positive cases with only a modest increase in false positives.

Precision–recall analysis demonstrated sustained robustness, with an AP of 0.928, comparable to the full-feature model. Predicted probability distributions showed clearer separation between CTS-positive and CTS-negative samples, with overlap largely confined to an intermediate-risk region (approximately 0.55–0.75). Calibration analysis demonstrated stable probabilistic behaviour across most probability bins, with close alignment to the identity line at higher predicted probabilities and increased variability in intermediate bins.

A key finding is that restricting the model to eight core variables can improve discrimination, operating-point performance, and class separation compared with a full-feature model comprising 34 variables. This improvement occurred despite a substantial reduction in input dimensionality, suggesting that most predictive information is concentrated within a small subset of clinically meaningful features. In routine clinical datasets, incompleteness, redundancy, and measurement variability are common. Consequently, additional variables may introduce noise rather than signal[2]. The improved performance of the simplified model supports the use of feature selection by importance over exhaustive inclusion of available measurements. It should be noted that the eight-variable subset is not intended to represent a globally optimal feature set, but rather a clinically interpretable and practically effective subset under the current dataset. Further optimisation with larger datasets is indispensable.

Similar to the full-feature model, permutation-based importance analysis confirmed that both ultrasound and electrophysiological measurements contributed to prediction within the simplified model. The most influential variable was the median nerve cross-sectional area at the pisiform level, followed by forearm calibre measurements and electrophysiological parameters including distal latency, conduction velocity, and potential amplitude. No single feature dominated performance, supporting the complementary role of structural and functional information.

### 3.4. Residual-based severity grading and structure– function mismatch

To better identify the heterogeneity within CTS severity grades, a residual-based grading framework was applied. Residual values spanned a wide range across the cohort. Left-sided residuals ranged from -38.4 to +24.7 m/s (mean 5.1 ± 13.4 m/s, median 8.8 m/s), while right-sided residuals ranged from -43.0 to +24.5 m/s (mean -4.4 ± 18.0 m/s, median -1.5 m/s). This wide dispersion indicates substantial variability in functional impairment not explained by ultrasound-derived morphology alone.

Within main severity grades, multiple residual-based subgrades were defined. For left wrists, grade 3 cases were distributed across subgrades 3a (n = 7), 3b (n = 6), and 3c (n = 7), while grade 2 cases spanned subgrades 2a (n= 1), 2b (n = 2), and 2c (n = 1). For right wrists, grade 3 cases similarly exhibited remarked heterogeneity, with subgrades of 3a, 3b, and 3c (n = 9 each). Lower severity grades also demonstrated residual-based subdivision, including subgrade 1b (n = 2) on the right side.

Residual-based subgrading provides additional insight into this decoupling. Within the same main severity grades, wide dispersion in residuals was observed. This reflected variability in functional impairment relative to morphology. Some patients exhibited electrophysiological deficits that exceeded those predicted from ultrasound morphology, while others showed significant morphological changes with relatively preserved nerve function. These patterns were present across severity levels, rather than being confined to advanced disease. These findings suggested that structure–function mismatch is an intrinsic feature of CTS rather than a late-stage phenomenon.

### 3.5. Clinical impact, limitations, and future work

From a clinical perspective, our findings support an ultrasound-led triage. Imaging serves as the initial screening step and informs down stream electrophysiological test rather than replacing it. Although residual-based analyses in this study rely on retrospective electrophysiological data, they reveal patterns of structure–function consistency and mismatch that help identify which patients are most likely to benefit from further electrophysiological test. In future clinical workflows, patients with high-confidence ultrasound findings and low or high model-predicted functional risk may be less likely to require routine electrophysiology. In contrast, patients with ambiguous ultrasound findings and intermediate risk may benefit from further electrophysiological tests. Such a strategy has the potential to improve assessment and allocate electrophysiological resources more efficiently.

Our study demonstrates the feasibility of applying data-driven models to routine clinical data characterised by small sample size and substantial incompleteness. The use of random forests with surrogate splits and OOB evaluation enabled robust performance estimation without reliance on aggressive imputation or artificial data augmentation. Importantly, calibration analysis showed that predicted probabilities remained clinically interpretable, supporting their use for risk-based stratification rather than binary decision-making alone.

Several limitations should be acknowledged. Treating left and right wrists from the same patient as independent samples may introduce intra-subject correlation, which could potentially inflate performance estimates. Although this approach reflects clinical practice and increases sample size, bilateral measurements are not fully independent and may share common physiological factors. This was a single-centre retrospective study with a small sample size, which may limit generalisability. Although random forests with surrogate splits were selected for their robustness to missing and heterogeneous data, OOB evaluation alone may not fully capture generalisation performance. External validation in independent cohorts will be essential to confirm the robustness of the simplified model and residual-based grading framework. Electrophysiological measurements were treated as the functional reference standard, although they are subjected to variability and may not fully capture patient-reported symptoms or functional impairment. Ultrasound features were manually measured thus highly relied on the experience of the operators, which may introduce measurement heterogeneity. While the data pattern reflects real-world practice, it may also attenuate associations between morphology and function.

Future work should focus on expanding sample size, multi-center validation, and extension of residual-based grading to longitudinal data to assess progression and treatment response. Future analyses should address intra-subject dependence and validate the proposed framework under patient-level data partitioning. More rigorous validation strategies, such as patient-level cross-validation or external testing on independent cohorts, will be applied to further assess model stability and reproducibility. A simpler manual calculation method for core variables and a rapid severity grading scales will be proposed to enhance generalization across institutions and scenarios. The framework presented here has the potential to be applicable to other compressive neuropathies and musculoskeletal conditions.

## 4. Conclusion

In this study, we demonstrate that CTS severity can be stratified using a limited set of complementary ultrasound and electrophysiological features. A simplified and clinically motivated model can outperform a higher-dimensional full-feature model under real-world data limitations. Importantly, residual-based grading reveals substantial structure–function mismatch within main severity categories, highlighting heterogeneity that is not captured by standard grading schemes. These results support an ultrasound-led triage in CTS assessment, with electrophysiology targeted to patients most likely to benefit.

## Acknowledgements

This work was supported by the UCL–Shanghai Jiao Tong University Strategic Partner Fund 2025/26 (No. 536735).

## References

[1] Hannaford A, Vucic S, Kiernan MC, and Simon NG. Review article “spotlight on ultrasonography in the diagnosis of peripheral nerve disease: the evidence to date”. International Journal of General Medicine, 14:4579–4604, 2021. 4

[2] Lee CH and Yoon HJ. Medical big data: promise and challenges. kidney research and clinical practice. Kidney research and clinical practice, 36(1):3–11, 2017. 4

[3] Shi D, Diao X, Shi L, Tang H, Chi Y, Li C, and Xu H. Charformer: A glyph fusion based attentive framework for high-precision character image denoising. MM ‘22: Proceedings of the 30th ACM International Conference on Multimedia, pages 1147–1155, 2022. 2

[4] Atroshi I, Gummesson C, Johnsson R, Ornstein E, Ranstam J, and Roseń I. Prevalence of carpal tunnel syndrome in a general population. JAMA, 282(2):153–158, 1999. 1

[5] MacDermid JC and Wessel J. Clinical diagnosis of carpal tunnel syndrome: a systematic review. journal of hand therapy. Journal of Hand Therapy, 17(2):309–319, 2004. 1

[6] Fowler JR. Nerve conduction studies for carpal tunnel syndrome: gold standard or unnecessary evil? Clinical Neurophysiology Practice, 40(3):141–142, 2017. 2

[7] Padua L, Coraci D, Erra C, Pazzaglia C, Paolasso I, Loreti C, Caliandro P, and Hobson-Webb LD. Carpal tunnel syndrome: clinical features, diagnosis, and management. The Lancet Neurology, 15(12):1273–1284, 2016. 2

[8] Pelosi L, Arańyi Z, Beekman R, Bland J, Coraci D, HobsonWebb LD, Padua L, Podnar S, Simon N, van Alfen N, and Verhamme C. Expert consensus on the combined investigation of carpal tunnel syndrome with electrodiagnostic tests and neuromuscular ultrasound. Clinical Neurophysiology, 135:107–116, 2022. 2

[9] Padua L, Cuccagna C, Giovannini S, Coraci D, Pelosi L, Loreti C, Bernabei R, and Hobson-Webb LD. Carpal tunnel syndrome: updated evidence and new questions. The Lancet Neurology, 22(3):255–267, 2023. 2

[10] Shapiro LM, Kamal RN, Management of Carpal Tunnel Syndrome Work Group, and American Academy of Orthopaedic Surgeons. American academy of orthopaedic surgeons/assh clinical practice guideline summary management of carpal tunnel syndrome. JAAOS-Journal of the American Academy of Orthopaedic Surgeons, 33(7):e356–e366, 2025. 1

[11] Sonoo M, Menkes DL, Bland JD, and Burke D. Nerve conduction studies and emg in carpal tunnel syndrome: Do they add value? Clinical Neurophysiology Practice, 3:78–88, 2018. 2

[12] Murciano Casas MD, Rodríguez-Piñero M, Jimeńez Sarmiento AS, Álvarez López M, and Jimeńez Jurado G. Evaluation of ultrasound as diagnostic tool in patients with clinical features suggestive of carpal tunnel syndrome in comparison to nerve conduction studies: study protocol for a diagnostic testing study. PLoS One, 18(11): e0281221, 2023. 2

[13] Diao X, Shi D, Tang H, Shen Q, Li Y, Wu L, and Xu H. Rzcr: Zero-shot character recognition via radical-based reasoning. Proceedings of the Thirty-Second International Joint Conference on Artificial Intelligence Main Track, pages 654–662, 2023. 2

